# HapSolo: An optimization approach for removing secondary haplotigs during diploid genome assembly and scaffolding

**DOI:** 10.1101/2020.06.29.178848

**Authors:** Edwin A. Solares, Yuan Tao, Anthony D. Long, Brandon S. Gaut

## Abstract

**Background:** Despite marked recent improvements in long-read sequencing technology, the assembly of diploid genomes remains a difficult task. A major obstacle is distinguishing between alternative contigs that represent highly heterozygous regions. If primary and secondary contigs are not properly identified, the primary assembly will overrepresent both the size and complexity of the genome, which complicates downstream analysis such as scaffolding.

**Results:** Here we illustrate a new method, which we call HapSolo, that identifies secondary contigs and defines a primary assembly based on multiple pairwise contig alignment metrics. HapSolo evaluates candidate primary assemblies using BUSCO scores and then distinguishes among candidate assemblies using a cost function. The cost function can be defined by the user but by default considers the number of missing, duplicated and single BUSCO genes within the assembly. HapSolo performs hill climbing to minimize cost over thousands of candidate assemblies. We illustrate the performance of HapSolo on genome data from three species: the Chardonnay grape (*Vitis vinifera*), with a genome of 490Mb, a mosquito (*Anopheles funestus;* 200Mb) and the Thorny Skate (*Amblyraja radiata*; 2,650 Mb).

**Conclusions:** HapSolo rapidly identified candidate assemblies that yield improvements in assembly metrics, including decreased genome size and improved N50 scores. Contig N50 scores improved by 35%, 9% and 9% for Chardonnay, mosquito and the thorny skate, respectively, relative to unreduced primary assemblies. The benefits of HapSolo were amplified by down-stream analyses, which we illustrated by scaffolding with Hi-C data. We found, for example, that prior to the application of HapSolo, only 52% of the Chardonnay genome was captured in the largest 19 scaffolds, corresponding to the number of chromosomes. After the application of HapSolo, this value increased to ~84%. The improvements for mosquito scaffolding were similar to that of Chardonnay (from 61% to 86%), but even more pronounced for thorny skate. We compared the scaffolding results to assemblies that were based on another published method for identifying secondary contigs, with generally superior results for HapSolo.

## BACKGROUND

Traditionally, reference genomes have been produced from genetic materials that simplify assembly; for example, the first two plant species targeted for reference quality genomes, *Arabidopsis thaliana* [1] and rice (*Oryza sativa*) [2], were chosen in part because they naturally self-fertilize and are therefore highly homozygous. Other early genomes, such as those from *Caenorhabditis elegans* and *Drosophila melanogaster* [3, 4], were also based on inbred, highly homozygous materials. Recent sequencing of additional model and non-model species have continued to rely on near-homozygous materials, either through inbreeding [5, 6] or by focusing on haploid tissue [7, 8].

The reliance on homozygous materials is fading rapidly, however, for at least three reasons. The first is that it has become clear that inbred materials can misrepresent the natural state of genomes. A dramatic illustration of this fact is that some lines of maize purged 8% of their genome in only six generations of self-fertilization [9]; more generally, inbred genomes tend to be smaller than those based on outbreeding species [10, 11]. The second is that many species of interest cannot be easily manipulated into a homozygous state. Many animals fall into this category, such as mosquitoes [12], as do many perennial crops like grapes, which are highly heterozygous [13] and can be selfed but only with substantial fitness costs that limits homozygosity [14]. Finally, some important features and phenotypes - such as sex determination [15] and other important adaptations - can only be identified by analyzing heterozygous samples.

Fortunately, the resolution of highly heterozygous regions, which often contain large structural variants, is now possible due to improvements in sequencing technologies and their affordability. In theory, long-read sequencing technologies, like those from Pacific Biosciences and Oxford Nanopore, provide the capability to resolve distinct haplotypes in heterozygous regions, leading to the assembly of reference-quality diploid genomes [5, 16, 17]. Several genomes based on highly heterozygous materials have been published recently [13, 18–22], with many additional efforts ongoing.

Nevertheless, the assembly of heterozygous genomes still presents substantial challenges. One challenge is resolving distinct haplotypes in regions of high heterozygosity. Programs that assemble long-reads, such as FALCON and Canu [23], can fuse distinct haplotypes into the primary assembly. This haplotype-fusion produces genomes that are much larger than the expected genome size. When haplotypes are fused, either into the same contig or as different contigs into the primary assembly, the increased size and complexity of the assembly complicates down-stream approaches, such as scaffolding by Hi-C or optical mapping. In theory, Falcon-unzip [19] solves some problems by identifying alternative (or ‘secondary’) haplotigs that represent the second allele in a heterozygous region and then providing a primary assembly without secondary contigs.

It remains a difficult problem to identify and remove alternative contigs during assembly, but there are some suggested solutions. For example, Redundans identifies secondary contigs via similarities between contigs [24] and removes the shorter of two contigs that share some pre-defined level of similarity. Another approach, PurgeHaplotigs uses sequence coverage as a criterion to identify regions with two haplotypes [25]. The reasoning behind PurgeHaplotigs is that alternative alleles in a heterozygous region should have only half the raw sequence coverage of homozygous regions. Accordingly, the algorithm proceeds by first remapping raw reads to contigs, then flagging contigs with lower than expected read depth, and finally remapping and removing low-coverage contigs from the primary haplotype-fused assembly. A more recent approach, implemented in the purge_dups tool [26], builds on the coverage-based approach of PurgeHaplotigs. Purge_dups has been compared to PurgeHaplotigs and is superior based on a few exemplar assemblies [26].

Here we report another strategy, which we call HapSolo, to identify and remove potential secondary haplotigs. Our approach is similar to Redundans, in that it begins with an all-by-all pairwise alignment among contigs and uses features of sequence alignment as a basis to identify potential alternative haplotigs. However, HapSolo is unique in exploring the parameter space of alignment properties to optimize the primary assembly, using features of BUSCO scores as the optimization target. Here we detail the approach and implementation of HapSolo, demonstrate that it efficiently identifies primary vs. secondary haplotigs and show that it improves HiC-based scaffolding outcomes relative to purge_dups. HapSolo has been implemented in python and is freely available (https://github.com/esolares/HapSolo).

## APPROACH and IMPLEMENTATION

### Pre-processing

Our method begins with the set of contigs from genome assembly. In theory, HapSolo will work for any set of contigs from any assembler and from any sequencing type (i.e., short-read, long-read or merged assemblies). Given the set of contigs, the first steps are to size sort the contigs and then to perform an all-by-all pairwise alignment among all contigs (**Figure 1, steps 1 and 2**), using each contig as both a reference and a query. In theory, pre-processing alignments can be performed with any algorithm, with the HapSolo implementation supporting either BLAT [27, 28] or minimap2 [29].

**Figure 1:**
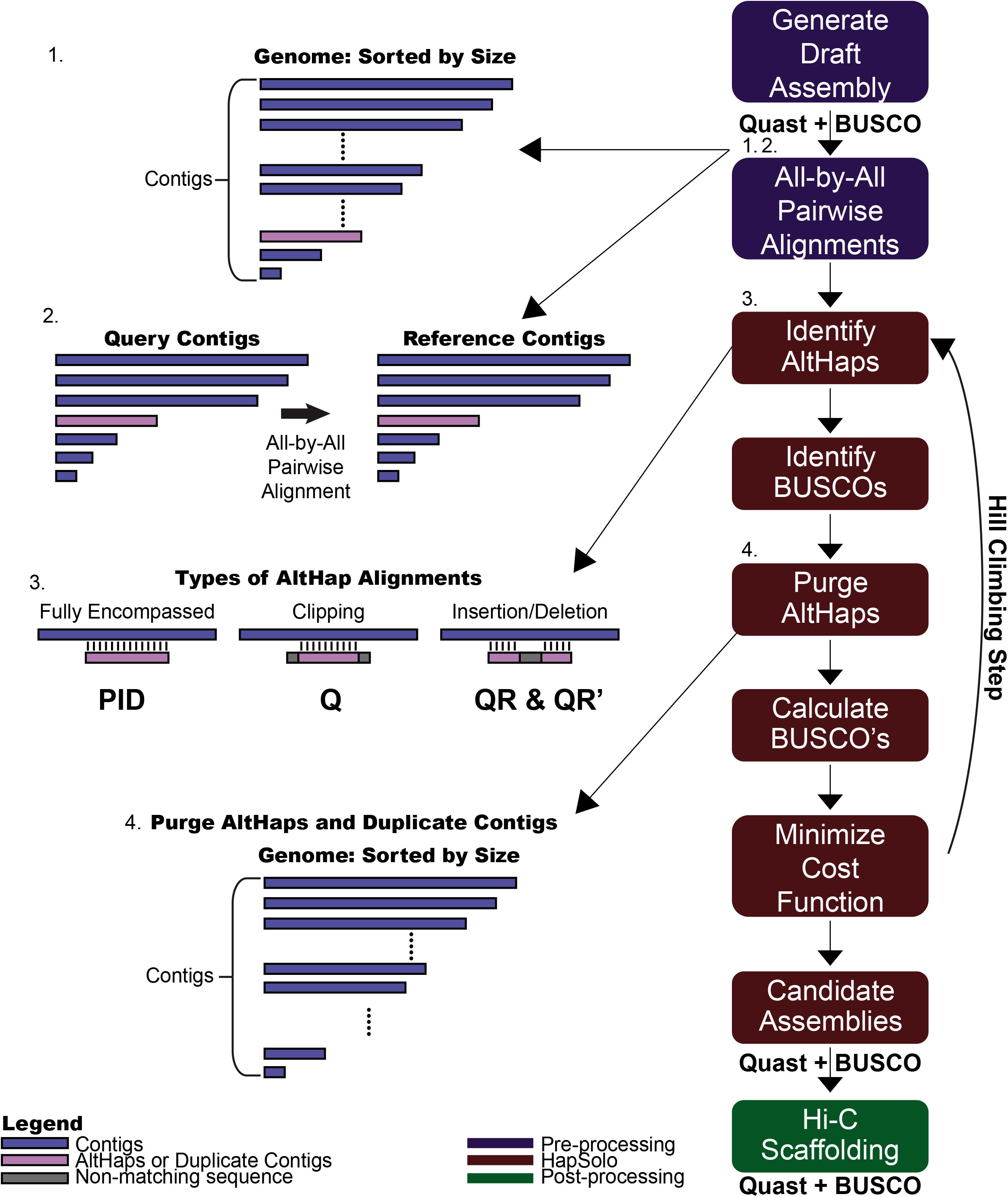
A schematic showing the basic workflow and ideas behind HapSolo. The rectangles on the right illustrate the basic steps, including pre-processing (blue rectangles), steps within HapSolo (red rectangles) and post-processing (green rectangle). Some of the HapSolo steps include iterations to perform hill climbing calculations, as described in the text and shown by the arrow. On the left, step 1 shows the contigs from the primary assembly, and step 2 illustrates the all-by-all alignment of contigs. Step 3 provides examples of some properties of potential alignments. The metrics - ID, Q and QR - were defined to help capture some of the variation among these conditions. Step 4 illustrates that new primary assemblies are formed by dropping putative secondary contigs.

### Steps within HapSolo

HapSolo imports alignment results into a PANDAS (https://pandas.pydata.org/) dataframe to form a table with rows representing pairs of aligned contigs and columns containing descriptive statistics for each pairwise comparison (**Figure 1, step 3**). Columns include the percent nucleotide identity between contigs (*ID*), a metric similar to those used in previous haplotig reduction programs; the proportion of the query contig length that aligns to the reference contig (*Q*), which is included to recognize that alignments can be clipped; and the ratio of the proportion of the query aligned to the reference relative to the proportion of the reference aligned to the query (*QR*). *QR* is considered because it reflects properties of aligned length and potential structural variant differences between contigs. A downside of *QR* is that it can reach values > 1.0, as longer variants may exist in either the query or the reference, and it is also non-symmetric. To compensate for this we include a symmetric value, which we define as *QR’=e*^*log*_2_(*QR*)^. The four parameters - *ID, Q, QR* and *QR’* - are the basis for filtering query contigs from the table and defining them as putative secondary contigs. For simplicity, however, we will emphasize *QR*, because *QR’* is dependent on *QR*.

In addition to the alignment table, HapSolo generates a table of BUSCO properties [30] for each contig. This BUSCO analysis is performed on each contig of the assembly prior to running HapSolo’s reduction algorithm. To perform these analyses, contigs are split into individual FAStA files and then BUSCO v3.0.2 is run on each contig separately so that they can be evaluated in parallel. Ultimately, the BUSCO table generated by HapSolo contains a list of complete (C) and fragmented (F) BUSCO genes for each contig. This table is integral for rapidly evaluating potential candidate assemblies.

Given the alignment table and the BUSCO table, HapSolo begins by assigning threshold values for *ID, Q* and *QR*, which we denote as *IDT, QT* and *QRT*. The threshold values can be assigned randomly, with set default values or with values defined by the user. The threshold values are applied to the alignment table to identify query contigs for purging. To be removed, a query contig must be in a pairwise alignment that satisfies three conditions: 1) an *ID* value ≥*ID_T_;* 2) e a *Q* value ≥*Q_T_*; and 3) a *QR* value that falls within the range min(*QR_T_,QR’_T_*) and max(*QR_T_,QR_T_*). After purging query contigs, HapSolo calculates the number of Fragmented (F), Missing (M), Duplicated (D) and Single-Copy (S) BUSCO genes across all of primary contigs that remain in the candidate assembly, based on values in the BUSCO table. It then calculates the *Cost* of the candidate assembly as:

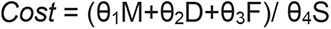

where θ_1_, θ_2_, θ_3_, and θ_4_ are weights that can vary between 0.0 and 1.0. Weights can be assigned by users; for all of our analyses below, we employ weights of 0.0 for F and 1.0 for M, D and S.

We then employ hill climbing to minimize *Cost* (Figure 1). Once *Cost* is calculated with random starting values, *ID_T_, Q_T_* and *QR_T_* are modified at each iteration by a randomized step size in the positive direction, which in turn defines a new set of primary contigs for a new cost evaluation. The steps consist of a fixed increment, which can be set by the user but is set to 0.0001 by default, multiplied by a random value sampled from U(0,1). As such, HapSolo utilizes a randomized forward walking agent to traverse the search space. If *Cost* does not change with new parameter values for a specified number of steps or if parameters increase past their maximum limits of *ID_T_ = Q_T_* = 1.00, then HapSolo assigns new random values of *ID_T_, Q_T_* and *QR_T_*. The process is repeated for *n* total iterations, and the iteration(s) with the smallest *Cost* are used to define the final set of primary contigs. When there are multiple solutions that minimize *Cost*, we retain all unique solutions; these additional solutions can be exported by the user for post-processing steps and evaluation. The values that determine the behavior of this minimization - e.g., the threshold for the number of consecutive cost plateaus, the number of *x* unique best candidate assemblies retained, the increase in step size by a fixed value, and the total number of iterations - can be set by the user.

To retain candidate assemblies with the smallest *Cost*, we implemented a unique priority queue (UPQ). The UPQ maintains a maximum number of *x* best assemblies, where *x* can be set by the user. The UPQ initially takes a list of one set of values, the score, primary contigs and other assembly information. The UPQ then takes the number of primary contigs for each of the candidate assemblies and sorts them by size. It then compares only the candidate assemblies of the same size, because assemblies of unequal size cannot be the same assembly. Therefore our algorithm, in order to reduce the number of contig set comparisons, only compares contig sets of the same size. Once it is established that the candidate assemblies of the same number of contigs are equal, only the candidate assembly with the lowest score is saved. The list is then sorted by score and returned. This allows retention of the max score of the best *x* number of assemblies by looking at the score of the last candidate assembly in the list, giving *O(1)* access to this value. Sorting takes *O(x log(x))*, where *x* is the best number of candidate assemblies to return, giving our UPQ a time complexity of *O(x log(x))*. Since we can instantaneously access the worst of the *x* candidate assemblies, we then perform an integer comparison of the score of our current candidate assembly with the worst score of our best *x* number of assemblies, reducing our computational time complexity. Only assemblies with the same or lower scores than the worst candidate assembly are then added to our UPQ. This reduces our total time complexity to *O(i x log(x))* where *i* is the number of iterations which produce scores lower than our max of *x*, and *x* is the number of best candidate assemblies to keep.

### Post-processing

Once HapSolo converges on a set (or *x* sets) of primary contigs that minimize *Cost*, the contig set is employed for post-processing steps to evaluate the candidate assembly. Specifically, we run QUAST v4.5 [31] and BUSCO 3.0.2 on the set of primary contigs that represent the best (or set of *x* best) candidate assemblies. QUAST measures basic genome assembly statistics, such as, N50, total assembly length, L50 and the largest contig size. Although not part of the HapSolo method, we provide scripts that run QUAST and BUSCO to output their results into a single score file.

### Implementation and requirements

HapSolo has been implemented and optimized for Python 2.7, but it is also supported under Python 3. However, we recommend using Python 2.7, for faster run times. HapSolo requires the input of a contig assembly (as a FAStA file), the location of a directory for individual contig BUSCO results, and the input of pairwise alignments. It currently supports either BLAT or minimap2 alignment output files (PSL or PAF or compressed PSL.gz or PAF.gz file).

## RESULTS & DISCUSSION

### Primary Assemblies

We illustrate the application and results of HapSolo on three diploid genome data sets. The three - including the Chardonnay grape (*Vitis vinifera*), the Anopheles mosquito (*A. funestus*) and the Thorny Skate (*Amblyraja radiata*) - represent a range of expected genome sizes, at 490Mb [32], 200Mb [20] and 2,560Mb (https://vgp.github.io/genomeark/Amblyraja_radiata/), respectively. The three datasets also represent a range of raw sequence coverage (at 58x, 240x, and 128x, respectively), and two different assembly methods - i.e., a hybrid assembly for Chardonnay [13] and Falcon_Unzip for both mosquito [20] and thorny skate [33]. The sequencing data are based on the Pacific Biosciences (PacBio) sequencing platform, but HapSolo should be applicable to any contig assembly drafted from any long-read assembler.

For pre-processing, we utilized pairwise alignments with BLAT and minimap2 for the Chardonnay and mosquito data. To limit run time, we applied BLAT to the Chardonnay and mosquito data without long contigs (> 10Mb) as queries, because we reasoned that >10Mb contigs are unlikely to represent alternative haplotigs (see Methods). These long contigs were included as references, however, so that they are represented in pairwise alignments. We used only minimap2 for the larger skate genome, due to prohibitively long run times with BLAT.

For each species, we applied HapSolo with and without hill climbing and compared the outcomes to the original unreduced assembly. **Table 1** provides assembly statistics, and it illustrates improvements from the unreduced assembly, to the assembly without hill climbing (-HC) based on default values, and finally to the assembly with hill climbing (+HC), which is based on random starting values and 50,000 iterations. Focusing on Chardonnay, for example, the primary contig genome size declined 13% from the unreduced assembly to the -HC assembly and another 5% from the -HC assembly to the +HC assembly. Not surprisingly, as genome size decreased, so did the number of contigs included in the assembly, which fell from 2,072 to 1,369 (-HC) to 1,155 (+HC). However, contig N50 increased by 35% from 1.066 Mb to 1.441 Mb. Similar results were achieved after applying HapSolo to contigs from mosquito and thorny skate (**Table 1**). For both assemblies, the number of contigs, L50 and genome size decreased, while the contig N50 improved by 9% for both mosquito and the thorny skate. We note, however, that hill climbing did not increase N50 for the mosquito assembly much beyond that achieved by applying HapSolo for one iteration with its default values, suggesting that the default values performed well by this measure with this dataset.

**Table 1:** Contig assembly statistics for three primary assemblies for each of three species.

| Species | Chardonnay |  |  | Mosquito |  |  | Thorny Skate<sup>4</sup> |  |  |
| --- | --- | --- | --- | --- | --- | --- | --- | --- | --- |
| Assembly Type | No HapSolo<sup>1</sup> | HapSolo -HC<sup>2</sup> | HapSolo +HC<sup>3</sup> | No HapSolo | HapSolo -HC2 | HapSolo +HC3 | No HapSolo | HapSolo -HC | HapSolo +HC |
| # of Contigs | 2,072 | 1,369 | 1,155 | 1,073 | 674 | 666 | 16,218 | 14,494 | 12,937 |
| Contig Assembly Size (Mb) | 655.2 | 569.3 | 539.0 | 212.0 | 200.4 | 200.0 | 3,229.4 | 3,147.8 | 3,031.3 |
| Largest Contig (Mb) | 11.6 | 11.6 | 11.6 | 7.6 | 7.6 | 7.6 | 3.4 | 3.4 | 3.4 |
| Contig N50 (Mb) | 1.1 | 1.3 | 1.4 | 0.6 | 0.7 | 0.7 | 0.4 | 0.4 | 0.5 |
| Contig L50 | 141 | 106 | 95 | 86 | 77 | 77 | 2,022 | 1,928 | 1,800 |

<sup>1</sup> Results in this column are based on the primary assembly without application of HapSolo
<sup>2</sup> Results in this column are based on application of HapSolo without hill climbing (-HC) and with default parameters of ID, Q and QR = 0.70.
<sup>3</sup> Results in this column are based on application of HapSolo with 50,000 cycles of hill climbing (+HC).
<sup>4</sup> Results for HapSolo were generated using minimap2. Chardonnay and mosquito statistics are based on BLAT.

Although N50 did not decline for the mosquito data, our implementation of hill climbing reduced *Cost*, as we expected, with the expected effects on BUSCO scores. **Figure 2** illustrates a sorted representation of *Cost*, showing that lower *Costs* were identified. The behavior of hill climbing is dependent on the assembly, starting values for the three parameters (*ID_T_, Q_T_* and *QR_T_*), and the number of local minima in the *Cost* function. Nonetheless, substantial improvements occurred within the first 1,000 iterations for all three datasets (**Supplementary Figure 1**), with only minor improvements thereafter. Overall, the improvement in *Cost* suggests value in applying hill climbing to new data sets, especially given that the computational costs are minor (see below).

**Figure 2:**
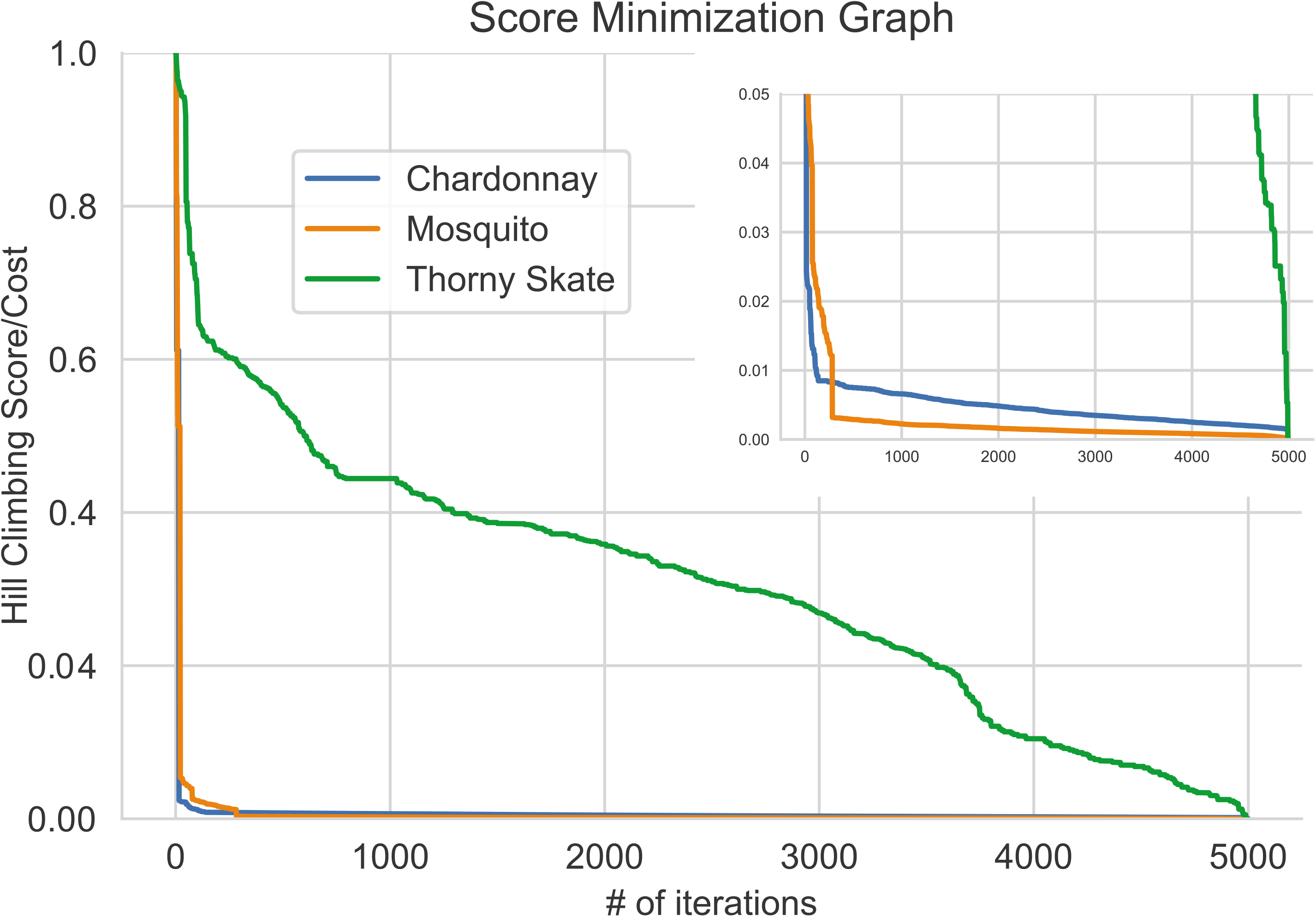
A graph of the sorted performance of hill climbing over 5,000 iterations, with normalized *Cost* on the *y*-axis and the number of iterations on the *x*-axis. For most of our analyses with HC, we performed 5,000 iterations on each of 10 cores; here we are showing results from one core. The top right provides a graph with altered scale for better visualization of Chardonnay and mosquito results.

**Table 2** complements information about *Cost* by reporting BUSCO scores. HapSolo achieved its principal goal, which is to generally increase the representation of single copy (S) BUSCO genes and decrease duplicated (D) genes in reduced compared to unreduced assemblies. Note the differences between the -HC and +HC assemblies, because in some cases the -HC assembly had more single copy genes but at the cost of also having more duplicated genes. Thus, the +HC option can produce assemblies with lower *Cost* but with fewer BUSCO genes.

**Table 2:** Starting and Ending BUSCO values for the three species for primary contig assemblies.

|  | No HapSolo |  | HapSolo (-HC) |  | HapSolo (+HC) |  |
| --- | --- | --- | --- | --- | --- | --- |
| Species | GS<sup>1</sup> | BUSCO<sup>2</sup> | GS | BUSCO | GS | BUSCO |
| **Chardonnay** | 655.2 | C:1357<br>S:1004<br>D:353 | 569.3 | C:1356<br>S:1152<br>D:204 | 539.0 | C:1357<br>S:1205<br>D:152 |
| **Mosquito** | 212.0 | C:2640<br>S:2493<br>D:147 | 200.4 | C:2609<br>S:2548<br>D:61 | 200.0 | C:2621<br>S:2566<br>D:55 |
| **Thorny Skate** | 3,229.4 | C:2091<br>S:1651<br>D:440 | 3,147.8 | C:2087<br>S:1675<br>D:412 | 3,031.3 | C:2080<br>S:1715<br>D:365 |

<sup>1</sup> Genome Size (GS) based on the sum of all contigs for the primary assembly.
<sup>2</sup> Busco based on all contigs prior to the application of HapSolo. The four values represent the complete (C), the single (S) and duplicated (D) BUSCO genes.

**Figure 3** plots the cumulative contig assembly length for the three assemblies for each of the three species, and it illustrates two important points. First, HapSolo reduced the total assembly length primarily by removing numerous contigs of small size. Second, differences between the -HC and +HC reduced assemblies were more evident for some species (e.g., thorny skate) than for others (e.g., Chardonnay). Nonetheless, when there were differences, hill climbing decreased both assembly size (**Table 1**) and *Cost* (**Table 2**).

**Figure 3:**
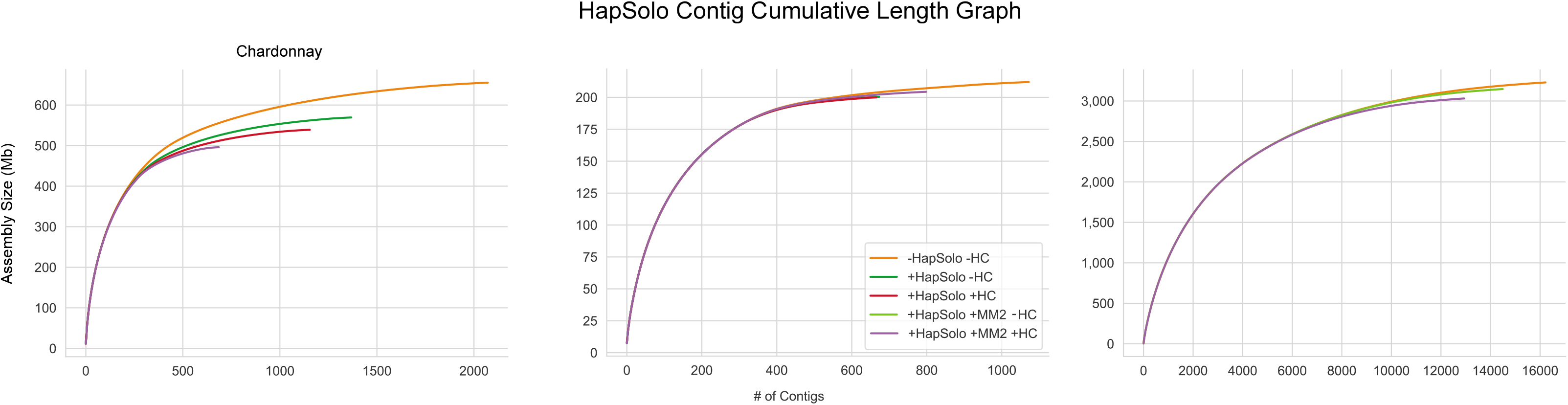
The cumulative assembly size (cdf) based on contigs. For each Chardonnay and mosqutio, different reduction strategies are depicted: an unreduced assembly, HapSolo applied with default parameter values and no hill climbing (-HC) using BLAT or minima2 (MM2), and HapSolo with random starting values and 50,000 iterations of hill climbing (+HC) using BLAT or minimap2 (MM2). For thorny skate we only have minimap2 based purging.

### Hi-C Scaffolding Results

HapSolo focuses on the improvement of primary assemblies, but there are potential advantages for removing haplotigs for downstream operations like scaffolding. Failing to remove duplicate haplotigs can cause false joins between duplicate haplotigs or lead to non-parsimonious joins between duplicate haplotigs and adjacent single copy regions. Here we illustrate the advantage of running HapSolo on primary assemblies prior to Hi-C scaffolding. For these analyses, the unreduced assembly and both reduced assemblies (i.e., -HC and +HC) were scaffolded using the 3D-DNA pipeline [34], resulting in more contiguous assemblies overall. We compared the improvements of the two scaffolded HapSolo assemblies against the unreduced scaffolded assembly (**Table 3**). Gains in improvements to the largest scaffold were clear across all assemblies relative to the unreduced assembly. For example, the largest scaffold increased by 1.71x (-HC) and 1.91x (+HC) for Chardonnay and by 1.22x (-HC) and 2.18x (+HC) for mosquito (**Table 3**).

**Table 3:** Scaffolded assembly statistics after Hi-C analysis on HapSolo assemblies, for three primary assemblies for each of three species.

| Species | Chardonnay |  |  | Mosquito |  |  | Thorny Skate<sup>6</sup> |  |  |
| --- | --- | --- | --- | --- | --- | --- | --- | --- | --- |
| Assembly Type | No HapSolo<sup>1</sup> | HapSolo -HC<sup>2</sup> | HapSolo +HC<sup>3</sup> | No HapSolo | HapSolo -HC | HapSolo +HC | No HapSolo | HapSolo -HC | HapSolo +HC |
| # of Scaffolds | 1,332 | 2,748 | 2,403 | 1,211 | 603 | 611 | 14,238 | 12,269 | 10,009 |
| % of Genome in *k* largest scaffolds<sup>4,5</sup> | 52.0% | 89.0% | 84.0% | 61.9% | 68.3% | 85.7% | 31.5% | 104.7% | 106.3% |
| % of Genome in Scaffolds > 10Mb<sup>5</sup> | 44.0% | 94.0% | 94.0% | 93.5% | 94.7% | 94.1% | 40.4% | 103.3% | 105.5% |
| Scaffold Assembly Size (Mb) | 656.1 | 570.4 | 540.1 | 212.5 | 200.8 | 200.3 | 3,240 | 3,158 | 3,039 |
| Largest Scaffold (Mb) | 19.1 | 32.6 | 36.5 | 43.6 | 53.0 | 95.0 | 170.0 | 208.4 | 250.5 |
| Scaffold N50 (Mb) | 7.2 | 23.5 | 20.7 | 37.9 | 41.5 | 41.6 | 62.1 | 65.5 | 69.8 |
| Scaffold L50 | 28 | 11 | 11 | 3 | 3 | 2 | 16 | 35 | 13 |

<sup>1</sup> Results in this column were based on the primary assembly without application of HapSolo
<sup>2</sup> Results in this column were based on application of HapSolo without hill climbing (-HC) and default parameters of ID, Q and QR.
<sup>3</sup> Results in this column were based on application of HapSolo with 50,000 cycles of hill climbing (+HC).
<sup>4</sup> Percentage of genome in the largest *k* scaffolds, where *k* is equal to the number of chromosomes expected for each species.
<sup>5</sup> Percentages normalized using expected genome sizes of 490Mb, 200Mb and 2,560Mb for chardonnay, mosquito and thorny skate respectively.
<sup>6</sup> Results for HapSolo were generated using minimap2.

**Figure 4** illustrates the distribution of scaffolds for each of the three species under various HapSolo implementations. For each scaffold we measured the proportion of the genome that was contained in the *k* largest scaffolds, where *k* is the haploid number of chromosomes for each species. For example, Chardonnay has 19 chromosomes, and the 19 largest scaffolds based on the unreduced assembly represented 52% of the genome size. Following HapSolo haplotig reduction, the largest 19 scaffolds encompassed up to 93% of the total expected genome size of 490Mb. Similar improvements were identified for the two other species, with mosquito improving from 61.9% to 85.7% and thorny skate from 31.5% to 106.3%. The observation of 106.3% of the thorny skate being contained in the largest *k* scaffolds indicates that the expected genome size is incorrect or that there is a need for additional purging of haplotigs.

**Figure 4:**
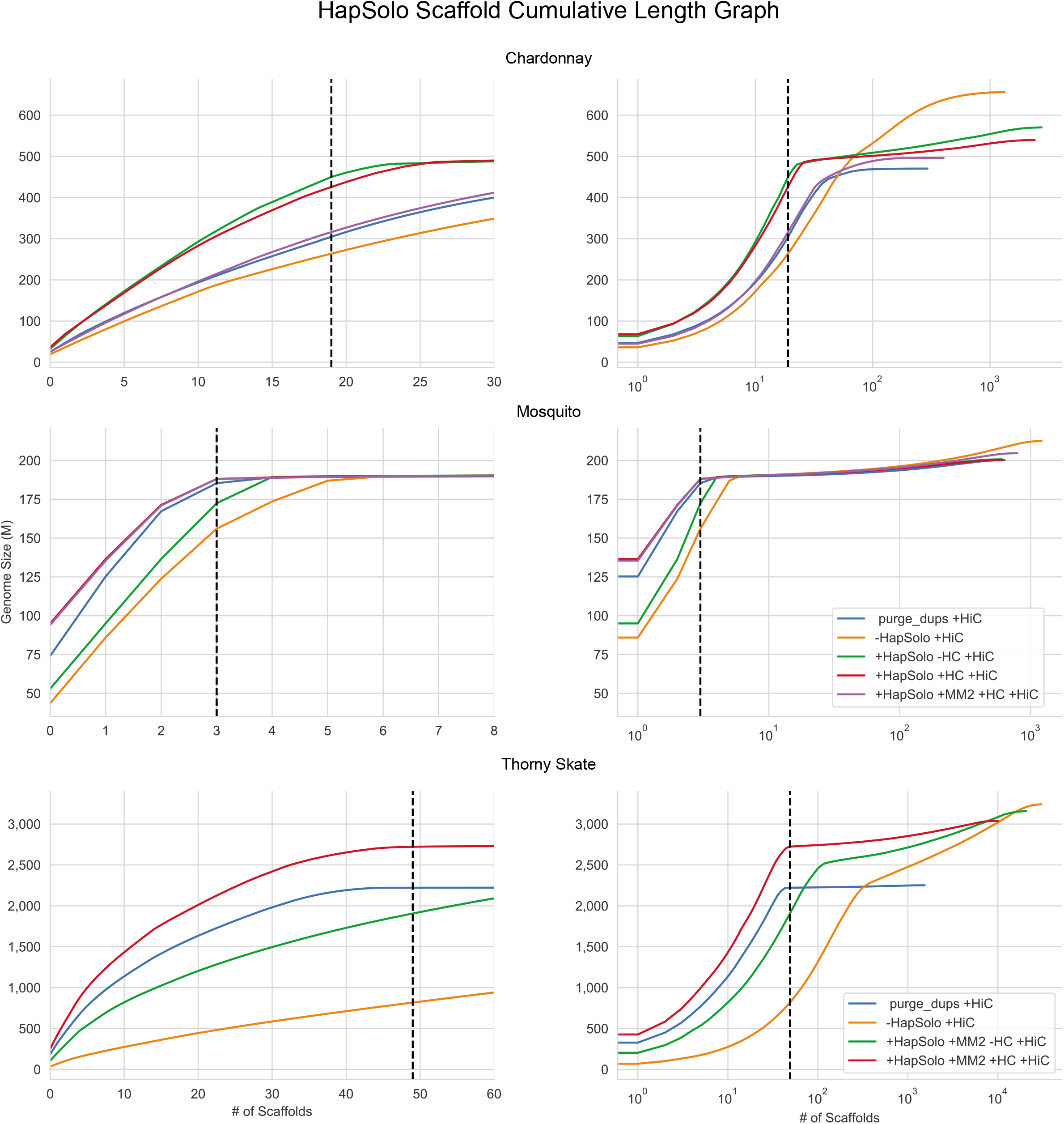
The cumulative assembly size (cdf) based on scaffolds for a different species in each row. There are two graphs for each species; the one on the left focuses on the chromosome length scaffold portion of the assembly (number of scaffolds), while the one on the right is the complete assembly on a log10 (number of scaffolds) scale. Each graph has five lines, representing the scaffolded genome based on purge_dups (purge_dups +HiC), the unreduced assembly (-HapSolo -HC +HiC), scaffold based on the HapSolo reduced assembly without hill climbing (+HapSolo -HC +HiC), and the scaffold based on the HapSolo reduced assembly with hill climbing (+HapSolo +HC +HiC), HapSolo reduced assembly with hill climbing using minimap2 (+HapSolo +MM2 +HC +HiC. In all graphs, the dotted line indicates the estimated number of chromosomes for the species.

HapSolo scaffolded assemblies were always demonstrably superior to the unreduced scaffolded assemblies for all three species, but the additional value of hill climbing varied among datasets.The value of hill climbing was clear for the mosquito, where the first 3 scaffolds (representing *k*=3 chromosomes) represented ~68% of genome with scaffolded -HC assembly *versus* 86% for the +HC reduced assembly. In contrast, hill climbing produced a disadvantage for Chardonnay (*k*=19, 92.6% -HC vs. 88.0% +HC) and only a small improvement for thorny skate (*k*=49, 104.7% -HC vs. 106.3% +HC). This being said, our metric based on the proportion of the genome in the *k* largest scaffolds is imperfect. For example, something as simple as a single split chromosome representing two metacentric arms could have a large effect on the metric. We therefore also examined other metrics, like the percentage of the genome encompassed in > 10Mb scaffolds and the longest scaffold. The largest differences were again due to application of HapSolo, with relatively minor differences associated with hill climbing (**Table 3**).

Finally, we focused on results based on comparing the two pre-processing alignment algorithms, BLAT and minimap2. We applied both algorithms to Chardonnay and mosquito. For mosquito, the results were similar with either aligner, but the BLAT results were markedly superior for Chardonnay (**Figures 3 & 4**). We do not know the cause of the discrepancy with Chardonnay, but we note that it is a genome that contains extensive structural variation between haplotypes, such that ~15% of genes are estimated to be in a hemizygous state [13]. We suspect that minimap2 often failed to extend alignments beyond large insertion and deletion events, even though we applied it with low gap and extension penalties substantially (see Methods). Minimap2 is, however, highly preferable for run times, and it can be applied easily to gigabase-scale genomes like thorny skate.

### Comparing HapSolo to an alternative method

Other algorithms have been devised to identify and remove alternative haplotigs [24, 26, 32]. In the publication of purge_dups, Guan et al. (2020) compared its performance to PurgeHaplotigs and found it to be generally superior. We compared HapSolo to purge_dups [26], focusing on scaffolding results after HiC analysis. **Figure 4** indicates that HapSolo generally led to better scaffolded assemblies than purge_dups, but with some caveats. For example, the HapSolo-based Chardonnay assembly was superior to the purge_dups assembly when BLAT was used to perform pre-processing. In this case, the percentage of the genome with >10Mb scaffolds was 97.7% for HapSolo versus 67.4% purge_dups, with a 32% improvement in largest scaffold (**Table 4**). However, purge_dups performed similarly to HapSolo for Chardonnay when pairwise alignments were based on minimap2 (**Figure 4**). For mosquito, purge_dups performed similarly to HapSolo with either preprocessing aligner, as long as hill climbing was included in HapSolo analysis. Finally, for the larger thorny skate genome, HapSolo with hill climbing outperformed purge_dups (**Figure 4**), resulting in a higher proportion of genomes in *k* scaffolds, more large (> 10Mb) scaffolds, and a 26% larger ‘largest scaffold’ (**Table 4**). Overall, HapSolo performed as well or better than purge_dups, based on the three exemplar datasets.

**Table 4:** A comparison of scaffolded assemblies after Hi-C analysis, based on HapSolo and purge_dups primary assemblies.

| Species<sup>1</sup> | Chardonnay |  | Mosquito |  | Thorny Skate |  |
| --- | --- | --- | --- | --- | --- | --- |
| Assembly Type | HapSolo +HC<sup>2</sup> | purge_dups | HapSolo +HC | purge_dups | HapSolo +HC<sup>5</sup> | purge_dups |
| # of Scaffolds | 2,403 | 294 | 611 | 635 | 10,009 | 1,534 |
| % of Genome in *k* largest scaffolds<sup>3</sup> | 88.0% | 58.5% | 85.7% | 83.7% | 106.3% | 86.8% |
| % of Genome in Scaffolds > 10M<sup>4</sup> | 97.7% | 67.4% | 94.1% | 92.7% | 105.6% | 86.0% |
| Scaffold Assembly Size (Mb) | 540.1 | 470.4 | 200.4 | 200.3 | 3,039.3 | 2,251.4 |
| Largest Scaffold (Mb) | 36.5 | 24.9 | 95.0 | 74.1 | 250.5 | 184.1 |
| Scaffold N50 (Mb) | 20.7 | 12.7 | 41.6 | 51.2 | 69.8 | 61.7 |
| Scaffold L50 | 11 | 15 | 2 | 2 | 13 | 11 |

<sup>1</sup> Data for HapSolo are based on BLAT alignments for Chardonnay and mosquito, and minimap2 alignments for thorny skate.
<sup>2</sup> Results in the +HC columns are based on application of HapSolo with 50,000 cycles of hill climbing (+HC).
<sup>3</sup> Percentage of genome in the largest *k* scaffolds, where *k* is equal to the number of chromosomes expected for each species. Percentages are normalized using expected genome sizes of 490Mb, 200Mb and 2,560Mb for Chardonnay, mosquito and thorny skate respectively.
<sup>4</sup> Percentages normalized using expected genome sizes of 490Mb, 200Mb and 2,560Mb for chardonnay, mosquito and thorny skate respectively.
<sup>5</sup> Results for HapSolo were generated using minimap2.

### Applying HapSolo to a genome with low heterozygosity

HapSolo was designed to address a specific problem: the assembly of highly heterozygous genomes with divergent haplotigs. We chose our three exemplars to represent the problem. But how does HapSolo perform on less heterozygous genomes? We applied HapSolo to the mouse, *Peromyscus leucopus*, a mammalian genome from a single diploid individual with low (0.33%) heterozygosity [35]. In samples with low heterozygosity, alternative haplotigs are less likely to exist, and hence we expect fewer benefits with the application of HapSolo. Indeed, we found no benefit. Comparing results between the reference assembly and the HapSolo assembly (with minimap 2 and hill climbing), we found similar proportions of the genome encompassed in 10Mb scaffolds (96.8% vs. 95.7%) and a substantially smaller proportion of the genome encompassed in *k*=24 chromosomes (88.5% vs. 71.1%) (**Table S1**). The HapSolo assembly was, however, largely contiguous with the reference assembly (**Figure S2**). Interestingly, purge_dups did not find any alternative contigs on this assembly and ultimately failed with an error, so we are unable to compare its performance.

In addition to the low heterozygosity, the *P. leucopus* genome has a low percentage of duplicated BUSCOs relative to the complete set of BUSCOs, at 2.1% (**Table S1**). In contrast, Chardonnay, mosquito and thorny skate have 26.0%, 5.5% and 21.0%, respectively (**Table 2**). Perhaps unsurprisingly, given this statistic, mosquito exhibits the least dramatic improvements in assembly statistics after application of HapSolo (**Table 3**). These observations suggest that there are lower limits at which HapSolo becomes ineffective and perhaps even detrimental. Based on the data we have analyzed, we suggest that ~5% may be a lower limit for the proportion of duplicated BUSCOs. Heterozygosity is likely to define another lower limit. Given that heterozygosity is 0.33% for *P. leucopus*, we expect that HapSolo will not be useful for the assembly of human genomes, because species-wide human heterozygosity is 0.05% [36]. Our results nonetheless suggest that HapSolo is likely to be a helpful tool for assemblies with a high number of duplicate BUSCOs.

### Execution Time and Memory Efficiency

To measure runtime, HapSolo was run on dual CPU Intel E5-2696 V2 nodes containing 512GB of RAM and storage attached via a 40Gbe connection. CPU runtime depends on the number of iterations, but it is also dependent on the data and parameter values. We measured runtime across the datasets, measuring different configurations in terms of the number of cores and the number of iterations per core (**Table S2**). Under the conditions we used for empirical data (i.e., hill climbing on 10 cores with 5000 iterations per core), the total time was <10 minutes for both Chardonnay and mosquito but substantially longer at 13 hours and 45 minutes for the much larger thorny skate. Note that memory usage was dependent on the size of the alignment file and independent of the number of iterations, because HapSolo stores alignments in memory for rapid filtering at each step during hill climbing. Nonetheless, the memory and speed requirements are such that HapSolo can be run on a laptop or desktop computer.

### Conclusions

We have presented an implementation, HapSolo, that is focused on improving primary assemblies by removing alternative haplotigs. In theory, the HapSolo package can be applied to any set of contigs from any assembly algorithm. The approach implemented in HapSolo is intended to replace laborious manual curation [37], and it follows some of the logic of existing programs, like Redundans [24], PurgeHaplotigs [25] and purge_dups [26]. However, HapSolo differs from competing programs by at least three features. First, it utilizes multiple alignment metrics, so that it is not reliant only on percent identity (*ID*). The goal of these multiple metrics is to better discriminate among some situations that may yield high identity scores but nevertheless lead to the retention of different contigs in the primary assembly (**Figure 1**). Second, when the hill climbing option is utilized, HapSolo relies on a maximization scheme based on BUSCO values. The underlying assumption is that maximizing the number of single-copy BUSCOs establishes more complete and less repetitive genomes. We emphasize that this is an assumption common to the genomics community, because most new genomes are reported with BUSCO scores to reflect their completeness and quality. Third, an important feature of HapSolo is the ability to modify the *Cost* function, so that the user may choose to weigh duplicated BUSCO genes less heavily or perhaps even ignore them altogether. This flexibility may prove useful for some applications. For example, it may be useful to ignore costs related to duplicated BUSCO genes when assembling polyploid genomes and instead focus only on complete and fragmented genes.

We have illustrated some of the performance features of HapSolo by applying it to data from three species that differ in genome size and complexity: Chardonnay grape, a mosquito, and the thorny skate. The common feature of these species is that diploid assembly is necessary. For all three species, we compared the unreduced primary assembly to two HapSolo assemblies, one that used default values (-HC) and one that used hill climbing minimization (+HC). Both HapSolo assemblies reduced genome size and markedly improved standard statistics like N50 (**Table 1** and **Figure 3**). The +HC contig assembly was generally better than the -HC assembly, but not always; the most substantial differences occurred between the unreduced assembly and either of the two HapSolo assemblies.

Our reduced assemblies scaffolded faster than unreduced assemblies and also led to more contiguous genomes. For each of our three species, the cumulative genome length associated with first *k* scaffolds (where *k* is the chromosome number) was much larger based on reduced vs. unreduced assemblies. The percentage of the genome contained in chromosome length scaffolds increased by at least 25% (**Table 3**). We conclude that in highly heterozygous samples that potentially have a large number of alternative haplotigs, some reduction step is critical for curating a primary assembly and for downstream scaffolding. This is true even when the primary assemblies are from Falcon_Unzip [19] which has already (in theory but perhaps not always in practice) identified secondary haplotigs. We further advocate for the use of the hill climbing feature in HapSolo, because the computational cost is relatively small but the gains can be large (**Figure 3**). Finally, we find that BLAT tends to outperform minimap2 as the pre-processing aligner and advocate for its use. However, it can be time prohibitive on large genomes, and hence HapSolo includes support for minimap2.

Based on the data in this paper, HapSolo generally led to similar or better outcomes than purge_dups [26], another recently published method to identify and remove haplotigs. That is not to say, however, that HapSolo cannot be improved. We can see two obvious areas for future growth. The first is to consider coverage statistics, which represents a point of departure between our approach and that of both Purge Haplotigs and purge_dups. We predict, but do not yet know, that the inclusion of coverage with our existing alignment statistics could lead to more accurate inferences. A second area of improvement may be to implement alternative maximization algorithms, such as simulated annealing. Finally, it may also be possible to include additional features in the calculations of *Cost*. Our present reliance on BUSCOs has the advantages of speed and wide acceptance in the genomics community. However, depending on the initial assembly, it is likely that some contigs do not contain a BUSCO gene, are therefore not considered in *Cost* and do not form the approximation of threshold parameters (*ID_T_, Q_T_* and *QR_T_*). It is not yet clear what additional features could be included in the *Cost* function, but *k*-mer representation is one possibility.

## METHODS

### Species and Data

The data for the assemblies for *V. vinifera* (cultivar Chardonnay) [13], *A. funestus* (mosquito) [20], and *A. radiata* (thorny skate) [33] were downloaded from public databases (see Data Availability). As mentioned, the contig assemblies were based on PacBio data. The chromosome number for each species was found in various sources [20, 33, 38]. The *P. leucopus* data were published in [35].

### Pre-processing

For each genome, pre-processing prior to application of HapSolo consisted of all-by-all pairwise contig alignments, as described above. For this study, we used BLAT v35 [28] and minimap2 [29]. BLAT was run with default options after the reference was compressed into 2bit format, and it was run using each contig as a separate query to reduce run time. Although not technically a feature of HapSolo, our github release provides a script to run Blat v35 [28] using this parallel approach. After running on individual contigs, the resulting PSL files were concatenated into a single PSL file for input into HapSolo. Minimap2 was used to compare feasibility and results between aligners; it was employed with the options” -P -k19 -w2 -A1 -B2 -O1,6 -E2,1 -s200 -z200 -N50 --min-occ-floor=100”.

### Assemblies, Hi-C data and Scaffolding

HapSolo was applied to with with default parameters of 0.70 for ID_T_, Q_T_ and QR_T_; hill climbing started with random values of ID_T_, Q_T_ and QR_T_ and then minimized *Cost* using hill climbing over 50,000 iterations. In HapSolo, BUSCO is run in *geno* mode on each contig using the orthoDB9 datasets and the AUGUSTUS species option. BUSCO v.3.0.2 relies on BLAST v.2.2.31+, AUGUSTUS v3.3, and BRAKER v1.9.

We obtained short-read Hi-C data from online public databases for scaffolding [13, 20] (see Data Availability). The Hi-C sequencing data were mapped to their respective assemblies using BWA [28]. The scaffolding of raw assembly and HapSolo processed assemblies were processed with the 3D de novo assembly pipeline v180419 [39], available from https://github.com/theaidenlab/3d-dna/. We ran QUAST v4.5 [31] for our post processing example and to assess performance during program development. For Figures 2 and S1, the normalized value was calculated by first subtracting the minimum observed Cost min(Cost) from the observed Cost. The numerator [Cost-min(Cost)] was then divided by [max(Cost)-min(Cost)].

### Computational Resources and Processing

For runtime analyses, HapSolo was run on dual CPU Intel E5-2696 V2 Nodes containing 512GB of RAM. The Blat, minimap2 and BUSCO pre-processing steps were run on these same nodes, but also one the UC Irvine High Performance Computing Cluster, Extreme Science and Engineering Discovery Environment (XSEDE) [40], San Diego Supercomputer Center (SDSC) Comet [41] and Pittsburgh Supercomputing Center (PSC) Bridges [42] clusters.

## ACKNOWLEDGEMENTS

We thank A. Minio from the Cantù Lab at UC Davis for informing us of the haplotig issue in genome assemblies, R. Lathrop and J.J. Emerson at UC Irvine for comments, and R. Kuo from the Roslin Institute, University of Edinburgh for devising the name, HapSolo.

## DECLARATIONS

### Ethics approval and consent to participate

Not Applicable

### Consent for publication

Not Applicable

### Availability of data and materials

*Vitis vinifera* data: NCBI under the BioProject ID PRJNA550461.

*Anopheles funestus* data: NCBI under the BioProject ID PRJNA494870.

*Amblyraja radiata* data: Genbank ID GCA_010909765.1, https://vgp.github.io/genomeark/Amblyraja_radiata/

*Peromyscus leucopus* data: Genbank ID GCA_004664715.2 Software: https://github.com/esolares/HapSolo

Publication Version: https://github.com/esolares/HapSolo/releases/tag/v0.1

### Authors’ contributions

EAS conceived of the approach, wrote the package and analyzed primary assemblies; YT performed Hi-C scaffolding; ADL and BSG provided ideas and feedback on the approach; EAS, ADL and BSG wrote the paper.

