## Supplemental Tables and Figures for "HapSolo: An optimization approach for removing secondary haplotigs during diploid genome assembly and scaffolding"

**Supplementary Table 1:** Scaffolded assembly statistics after Hi-C analysis, for three primary assemblies of *Peromyscus leucopus.* Minimap2 was used for pairwise alignment during the pre-processing stage.

| Species | *Peromyscus leucopus* | | |
|---|---|---|---|
| Assembly Type | No HapSolo[1] | HapSolo -HC[2] | HapSolo +HC[3] |
| # of Scaffolds | 1,728 | 1,369 | 1,365 |
| % of Genome in 24 largest scaffolds[4] | 88.5% | 76.1% | 71.1% |
| % of Genome in Scaffolds > 10Mb | 96.8% | 95.6% | 95.7% |
| Scaffold Assembly Size (Mb) | 2,475.1 | 2,420.6 | 2,402.7 |
| Largest Scaffold (Mb) | 454.1 | 134.6 | 133.9 |
| Scaffold N50 (Mb) | 89.3 | 69.4 | 63.9 |
| Scaffold L50 | 8 | 14 | 15 |
| BUSCO's | C:3860 S:3779 D:81 | C:3869 S:3795 D:74 | C:3862 S:3785 D:77 |

[1] Results in this column were based on the primary assembly without application of HapSolo

[2] Results in this column were based on application of HapSolo without hill climbing (-HC) and default parameters of ID, Q and QR.

[3] Results in this column were based on application of HapSolo with 50,000 cycles of hill climbing (+HC).

[4] Percentage of genome in the largest *k* scaffolds, where *k* is equal to the number of chromosomes expected for each species.

**Supplementary Table 2:** Run time and performance statistics for hill climbing analyses, based on the three exemplar genomes.

| | Cores | Iterations | Total Iterations | Avg. Time (h:m:s)[1] | RAM (MB) |
|---|---|---|---|---|---|
| **Chardonnay** | 1 | 5,000 | 5,000 | 00:08:00 | 86.4 |
| | 10 | 5,000 | 50,000 | 00:09:16 | 93.2 |
| **Mosquito** | 1 | 5,000 | 5,000 | 00:03:18 | 160.0 |
| | 10 | 5,000 | 50,000 | 00:03:53 | 154.0 |
| **Thorny Skate** | 1 | 5,000 | 5,000 | 13:39:28 | 302.0 |
| | 10 | 5,000 | 50,000 | 13:45:40 | 321.0 |

[1] h = hour, m = minute, s = seconds

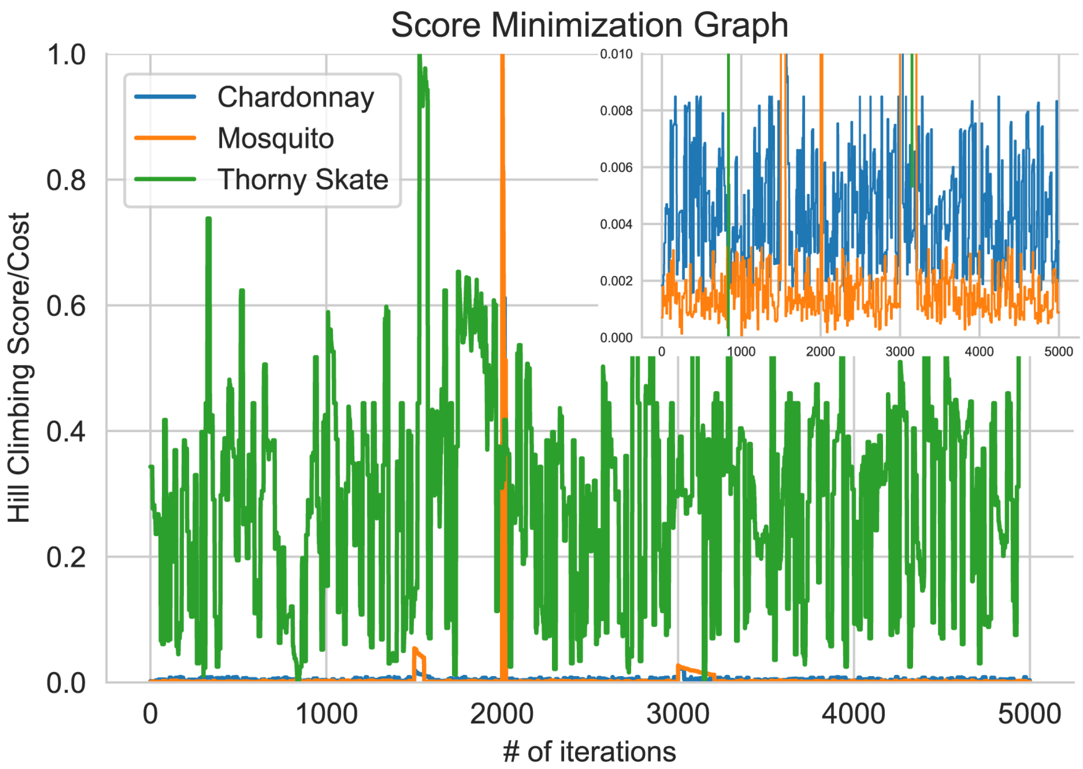


**Supplementary Figure 1:** A graph of the performance of hill climbing over 5,000 iterations, with normalized *Cost* on the y-axis and the number of iterations over time on the x-axis. The most dramatic improvements in *Cost* occurred within the first 1,000 iterations. For most of our analyses with HC, we performed 5,000 iterations on each of 10 cores; here we are showing results from one core. Costs were normalized by dividing the cost with the maximum cost observed for each respective assembly and subtracting the minimum observed cost to both values. In the top right, the zoomed-in version better illustrates behavior for Chardonnay and mosquito.

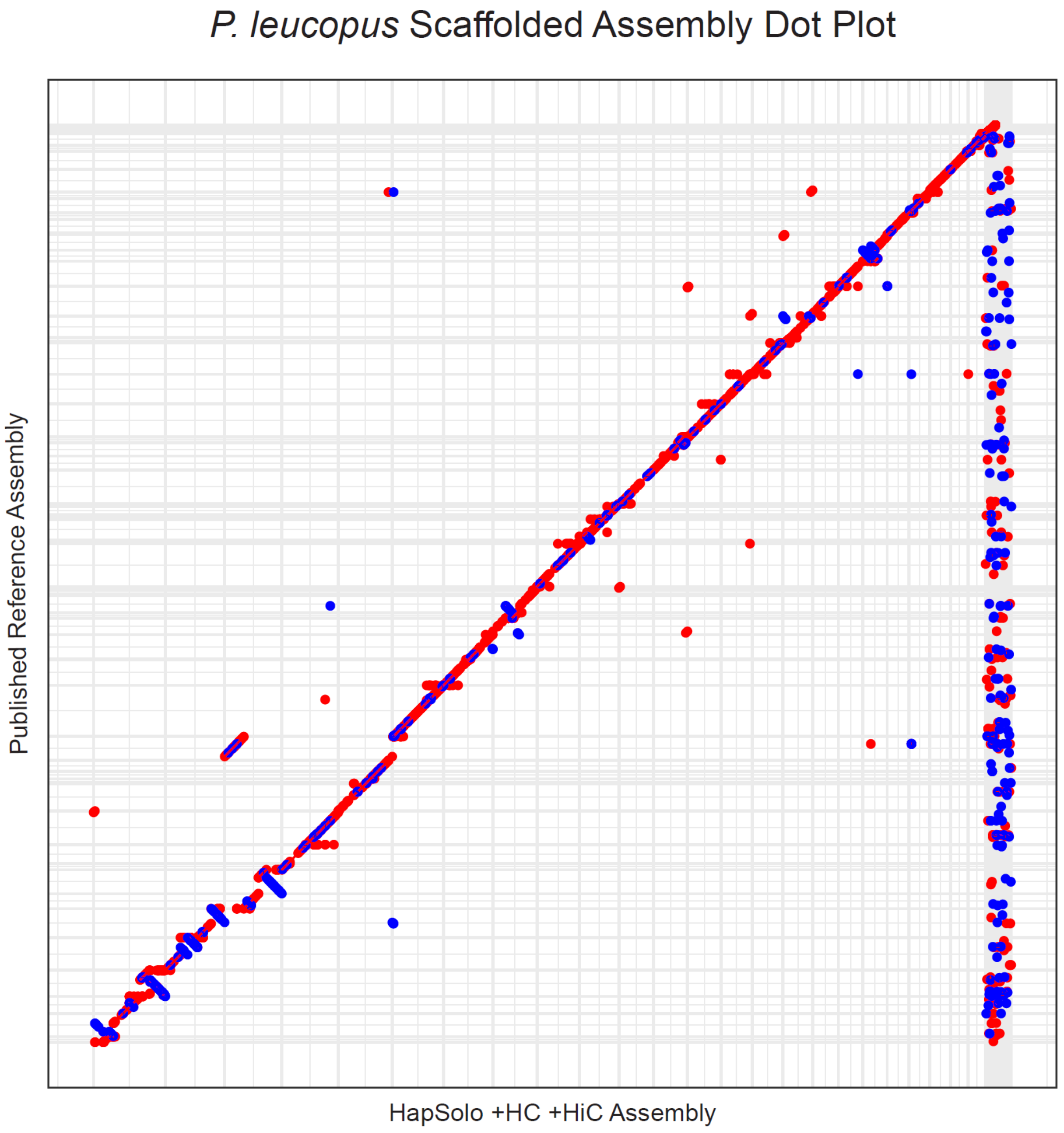


**Supplementary Figure 2:** A dot plot of *P. leucopus* assemblies. Red color dots denote sequences in the forward direction, and blue for sequences in the opposing direction. Grey lines illustrate boundaries for each scaffold. Blue or red colored dots that are found in multiple locations represent duplications. The dots forming a line decrease from left to right represent inversions or sorting errors.
